# Defending Synthetic DNA Orders Against Splitting-Based Obfuscation

**DOI:** 10.1101/2025.03.12.642526

**Authors:** Shaked Tayouri, Vladislav Kogan, Jacob Beal, Tal Levy, Dor Farbiash, Kevin Flyangolts, Steven T. Murphy, Tom Mitchell, Barak Rotblat, Isana Vaksler-Lublinsky, Rami Puzis

## Abstract

Biosecurity screening of synthetic DNA orders is a key defense against malicious actors and careless enthusiasts producing dangerous pathogens or toxins. It is important to evaluate biosecurity screening tools for potential vulnerabilities and to work responsibly with providers to ensure that vulnerabilities can be patched before being publicly disclosed. Here, we consider a class of potential vulnerabilities in which a DNA sequence is obfuscated by splitting it into two or more fragments that can be readily joined via routine biological mechanisms such as restriction enzyme digestion or splicing. We evaluated this potential vulnerability by developing a test set of obfuscated sequences based on controlled venoms, sharing these materials with the biosecurity screening community, and collecting test results from open source and commercial biosecurity screening tools, as well as a novel Gene Edit Distance algorithm specifically designed to be robust against splitting-based obfuscations.

## 1 Introduction

Over the past two decades, the engineering of biological organisms has been transformed from a difficult and costly research endeavor into a readily accessible engineering discipline [28]. Rapid development of biological systems is supported by large online libraries of genes [13,45,79], alongside a growing marketplace of nucleic acid synthesis companies that can quickly manufacture and ship genetic constructs for relatively low costs [16]. These constructs can be used to transform a wide variety of organisms by following various straightforward laboratory protocols [41, 46, 55].

The ubiquity and digitization of synthetic biology raise many security concerns in the emerging field of cyberbiosecurity [54]. Examples include threats to the security and privacy of genomic data and analysis [9, 17, 29, 35], vulnerabilities in biological laboratory instruments and automation [22, 56], biomedical software stack and supply chain security [24, 62, 67], and the potential for malicious actors to evade biosecurity screening in order to obtain dangerous pathogens or toxins [31, 48, 68].

Synthetic DNA poses a potential path by which malicious actors such as bioterrorists could acquire dangerous biological agents, such as venom components, toxins, or viruses [18, 47]. See § 2.1 for a brief background on DNA for non-biologists. Because of the dual use potential of synthetic biology for both benefit and harm [59], there is widespread agreement that biosecurity concerns should be taken into account in access to potentially dangerous DNA [31, 68, 72]. Thus, most synthetic DNA orders are evaluated for potential threats using biosecurity screening tools (§ 2.4), following guidelines (§ 2.2) from governments [14, 58], industrial consortia [19], and other organizations [38].

Here, we focus on a scenario where adversaries order dangerous DNA from a synthetic DNA provider while obfuscating their orders to circumvent screening. Similar to the obfuscation of image data [40, 53], DNA obfuscation is an adversarial input manipulation targeted to circumvent detection mechanisms. Unlike other adversarial input manipulations, the obfuscated DNA is later reconstructed in the physical (biological) world using a biological toolbox.

Biosecurity screening tools are not flawless, which makes it essential to identify and address potential vulnerabilities [48, 59]. In this paper, we explore a class of potential vulnerabilities that allows the obfuscation of DNA orders by splitting threat sequences into disjoint fragments (§ 3). The weakness of current DNA screening arises from the regulator’s requirement to flag malicious sequences longer than some predefined threshold (§ 2.2). Flagging short suspicious sequences dramatically increases the false positive rate. In § 4, we alleviate this gap using the Gene Edit Distance (GED) algorithm, which eliminates the need to define minimum detectable sequence length. Briefly, instead of checking if a DNA order contains a dangerous sequence, GED quantifies the effort required to transform the order into a dangerous sequence.

The efficacy of GED and state-of-the-art DNA screening tools at detecting obfuscated sequences were evaluated based on a carefully prepared benchmark dataset (§ 5). The contributions of this article can be summarized as follows:

- We introduce, validate, and evaluate the impact of two splitting-based DNA obfuscations on circumventing commercial and open-source biosecurity screening tools.
- Disclosure of the discovered weakness has led to the patching of a prominent open-source DNA screening tool (SeqScreen [4]).
- Obfuscated sequences produced in this research were included in a benchmark dataset produced by the International Gene Synthesis Consortium (IGSC).
- GED detects obfuscated sequences with a decent robustness margin.

Implications of this study, discussed in § 6, stress the necessity of continuous red teaming of synthetic biology pipelines in general and the biosecurity screening toolbox in particular. Further, the similarity of biosecurity and cybersecurity operations invites cybersecurity specialists to explore the biosecurity arena.

## 2 Background and related work

This section provides essential information for subsequent sections on potential cyberbiological threats and their mitigation. Readers familiar with protein synthesis and the enzyme restriction system are encouraged to skip § 2.1. Key terms are in **bold**.

### 2.1 Introduction to DNA editing

Genetic information in cells is encoded in DNA sequences of **nucleotides**, represented by cytosine (C), guanine (G), adenine (A), and thymine (T). DNA is a double-stranded molecule (**dsDNA**) with paired nucleobases (C-G, A-T) forming **base pairs (bp)**. Synthetic DNA costs $0.05 - $0.3 per bp and is typically delivered as cyclic dsDNA molecules called **plasmids**, which are stable and can replicate in living cell. Plasmids modify cell functionality through **transfection** processes.

DNA may affect cellular functionality in many ways. Here, we focus only on genetic instructions for building proteins (the actual toxins and venoms). Genes encoding proteins begin with **promoters** and end with **terminators**. Synthesizing a protein begins with creating a temporary copy of the DNA called **RNA**^1^ in a process called **transcription**. In computer terms, RNA can be compared to volatile memory, whereas DNA is considered persistent storage. Promoters determine where the transcription begins, which strand is transcribed, and in which direction. Promoters can be considered as function call pointers and terminators as the return commands.

Every three RNA nucleotides form a **codon**, which corresponds to one of the 20 natural **amino acids**. Multiple codons may be mapped to the same amino acid, and the choice of optimal codons varies between organisms. A process known as **translation** produces sequences of amino acids from RNA codon by codon, starting from the start codon. Short chains of amino acids are referred to as **peptides**, while longer chains form **proteins**. Both can be toxic.

An **intron** is a segment of a gene that does not code for proteins. During transcription, a copy of the gene’s entire sequence is made, including both introns and exons (the protein-coding regions). The cell then removes the introns between exons and joins the exons in a process called **splicing**. Introns are marked by specific sequences at their boundaries, known as splice sites, which signal where the splicing should occur. After splicing, the joined exons are translated into a protein. Splicing is the first among two biological tools we use for splitting-based DNA obfuscation.

The second tool that can be exploited for DNA obfuscation is the **restriction enzyme** system, which enables precise DNA editing in living organisms. This system relies on a specific enzyme and its corresponding **restriction site** sequence, with the enzyme acting as molecular scissors that cut the double-stranded DNA at precise locations called **cleavage** points. After cleavage, the loose ends of the DNA can be **ligated** by chance joining the disconnected ends and fixing the cleaved DNA. If multiple cuts are made by the same enzyme, the loose ends from different cuts can be ligated, creating a new sequence. In this article, an attacker exploits this common biological protocol to reconstruct obfuscated DNA.

### 2.2 Biosecurity regulations

Certain DNA sequences encode dangerous products, such as toxic peptides. Recognizing the threat, legislation tries to keep pace with technology. In 2020, California mandated all customers order synthetic genes from companies that perform gene screening [74]. In 2023 all government-funded researchers were recommended to purchase DNA only from companies that screen customers and orders [61]. Finally, starting in October 2026, orders will need to be screened against all “sequences known to contribute to pathogenicity or toxicity, even when not derived from or encoding regulated biological agents” [70, p. 6]. Today, synthetic DNA orders are screened for toxins and pathogens, including a large collection of regulated dangerous DNA and some sequences subject to export control.

The Screening Framework Guidance for Providers and Users of Synthetic Nucleic Acids (**HHS guidelines**), published by the United States Department of Health and Human Services, suggests methods to minimize the risk of unauthorized distribution of toxins and pathogens [58]. Closely related to the HHS guidelines are the Harmonized Screening Protocol v2.0 (HSPv2), used by the IGSC [19], and the International Association Synthetic Biology (IASB) Code of Conduct for Best Practices in Gene Synthesis [38]. The HHS guidelines, IASB Code of Conduct, and HSPv2 establish standards and practices aimed at preventing the misuse of synthetic genes. They define procedures for customer screening and synthetic gene order screening to detect possible toxins, pathogens, and other biological agents that pose a significant threat to public health and safety.

The HHS guidelines recommend screening for specific sequences, while the HSPv2 recommends identifying sequences derived from or encoding a toxin or pathogen. For example, a sequence alignment tool, such as BLAST, can compare gene orders with known sequences in the GenBank database [13]. The ***Best Match*** approach, as recommended by the HHS guidelines [58], involves searching every sequence **segment** of **50 bp**^2^ in the synthetic gene order within a database. The classification of the most similar sequence in the database determines the legitimacy of the order. If the Best Match of any fragment is a regulated toxin or pathogen, the order is flagged for further investigation.

The 50 bp threshold was chosen to maintain a manageable number of false alerts as screening shorter sequences often yields ambiguous results.

The HHS guidelines do not specify a database to use for screening, but they suggest GenBank as an example of such a database. The lack of a shared standard for identification is a known problem leading to inconsistent screening between companies, false positives due to genes shared between pathogenic and non-pathogenic organisms, and increased cost of overall screening [48], and a current area of development [75].

### 2.3 Cyber vs. biosecurity operations

At cybersecurity operation centers, analysts struggle with an overwhelming volume of security alerts that must be processed every day, leading to alert fatigue and the potential neglect of critical threats [3, 57]. Alert correlations are highly efficient, on the one hand, allowing to reduce thresholds for individual alerts, and on the other hand, reducing the overall burden of false positives. Threat intelligence is widely used to improve situational awareness (64%), prioritize alerts, and optimize the attention of analysts (50%), among other use cases [15]. The complexity of modern cyber threats, which often involve sophisticated techniques and multi-stage attacks, requires analysts to continuously update their knowledge and skills [63]. Although missing an attack is an undesirable event, until recently, software (including security) vendors were not liable for potential misuse of their products [34, 36].

Biosecurity analysts face similar challenges in their daily screening of synthetic DNA orders [61]. In general, the design and operation of DNA screening tools today are similar to signature-based intrusion detection. Alerts triggered by suspicious orders require follow-up investigation by human analysts, including contacting the customer [51]. On the one hand, false positive alerts increase screening costs and alert fatigue, just as with cybersecurity. On the other hand, false negatives may lead to dangerous misuse of the delivered products. There is currently a gap in alert correlation for biosecurity. The GED approach we propose in § 4 partially alleviates this gap by intelligently merging weak alerts raised within the same DNA sequence. The liability in such cases is still not well-defined [67]. Furthermore, although information sharing (i.e., threat intelligence) on suspicious cases is considered important [19, 58], it remains underused [32]. Overall, Diggans and Leproust [24] emphasize the importance of applying cybersecurity methods in the domain of biosecurity.

### 2.4 Biosecurity screening tools

**SeqScreen** [4] is a bioinformatics tool originally developed with funding from the IARPA Fun GCAT program, which focused on advancing biosecurity capabilities for sensitive and accurate analysis of short DNA sequences. SeqScreen attempts to report protein function with a special focus on functions of potential sequences of concern. The tool is regularly updated with new versions to ensure it incorporates the latest advancements and features for efficient pathogen detection.

**ToxinPred** [64] is a well-known tool for predicting peptide toxicity. The latest version, ToxinPred 3.0, showcases improved reliability and accuracy by incorporating machine learning and deep learning models.

Additionally, there are biosecurity screening algorithms, such as GenoTHREAT [1], BlackWatch [43], and NNTox [39], as well as various machine learning approaches [27,42,49,80] that can be used to predict the function of DNA and protein sequences. These approaches provide detailed information for human analysts to investigate a hit, including both suspicious and legitimate elements found within the DNA.

In response to growing biosecurity challenges, non-profit initiatives such as SecureDNA^3^ and IBBIS^4^ have introduced free screening tools designed to mitigate risks in DNA synthesis. SecureDNA is a screening platform that employs cryptographic technology to safeguard DNA synthesis [11]. While all code for system operation is open-source and accessible, the database and its generation process remain closed for security reasons. Similarly, IBBIS has developed the Common Mechanism, a free and open-source tool for DNA and RNA synthesis screening, including its databases [76]. These initiatives aim to establish a globally accessible baseline for preventing the exploitation of synthesis technology, supporting providers in effectively screening orders while addressing the challenges of high development costs and increasing economic burdens in the biotechnology sector.

A number of commercial organizations have also developed robust screening solutions. Companies such as Aclid^5^, RTX BBN Technologies [12, 78], and BATTELLE [30] maintain proprietary databases and tools optimized for rapid comparison of massive amounts of DNA data. Unlike academic frameworks, commercial tools must strictly adhere to the regulations, ensuring a comprehensive and compliant approach to biosecurity screening (see § 2.2).

## 3 Splitting-based DNA obfuscation

In the following subsections, we discuss the threat of an adversary obtaining DNA that produces components of dangerous biological agents (e.g., one of the toxins in a cone snail venom) despite security controls deployed by vendors to limit the dissemination of such DNA.

### 3.1 Threat model

#### Goal

The attackers assumed in this article could be a careless Do It Yourself (DIY) biologist, a frustrated student, a small startup company, etc., with goals ranging from pranks to designing new drugs in their garage. We will call them adversaries or attackers, although, in many cases, they may mean no harm but mistakenly or carelessly produce dangerous biological agents in their lab. The attacker would like to possess a dangerous biological agent (such as those often needed to produce drugs or vaccines) in an amount sufficient to harm a small group of people. Some neurotoxins are volatile and can potentially cause serious or lethal consequences. We assume the attacker does not aim to produce bioweapons, which would require sophisticated delivery methods. However, they do not have the means, the patience, or the knowledge to obtain the biosafety permits required for their experiments.

#### Attack flow

The general attack flow begins with designing the obfuscated malicious DNA (Figure 1 step 1). The attacker chooses a toxin, splits it into multiple fragments, and places the fragments within a larger DNA sequence (see § 3.3). The attacker orders the obfuscated DNA from a synthetic DNA provider (Figure 1 step 2). In this scenario, the packaging methods (e.g., standard plasmids, custom plasmid vectors, or retroviruses) of DNA molecules do not affect the interplay between attackers and defenders. For simplicity, we assume that the DNA is packaged within a standard plasmid. The order is screened by the provider (Figure 1 step 3). If it passes screening, the DNA is manufactured and delivered to the attacker after passing quality control procedures (Figure 1 steps 4-6). Finally, upon delivery, the attacker performs the biological procedures required to reconstruct the obfuscated DNA and produce the toxin/venom (Figure 1 step 7). Our study focuses specifically on steps 1, 3, and 7.

**Figure 1:**
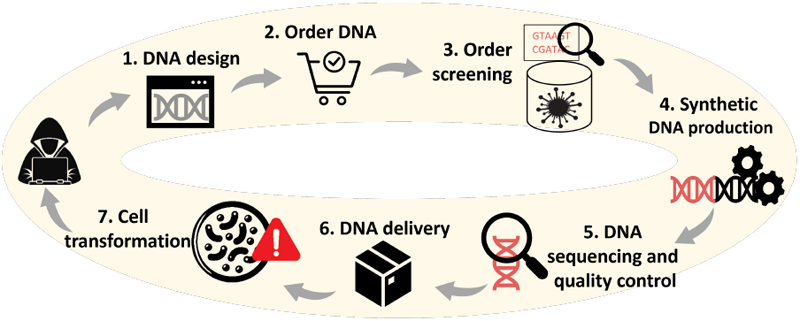
A typical synthetic biology workflow, and attack components.

#### Defense

Our analysis relies on standard biosecurity screening practices currently accepted within the synthetic DNA industry and the research community (Figure 1 step 3, § 2.2). We assume the attacker can only purchase DNA from providers that adhere to the HHS Guidelines. While this is not yet the case globally, it is reasonable to assume that the necessary regulations (§ 2.2) will soon come into force in most civil countries. Additionally, we assume that all orders are screened and the screening tools’ databases are sufficient to detect any toxin/venom sequences the attacker may attempt to acquire. Despite this, the attacker still attempts to bypass the screening mechanisms.

We also assume the attacker is familiar with the HHS Guidelines, has access to public sequence databases, BLAST, and all code libraries to implement the guidelines and test their sequences. However, the attacker has no access to the specific screening tools or proprietary databases employed by the synthetic DNA provider from which they intend to order the malicious DNA.

#### Attacker skills and resources

We assume an intermediate-level attacker, as defined by STIX Core Concepts [65], with skills and resources comparable to an average individual. Although the attackers’ skill level required to circumvent biosecurity screening is intermediate, it is multi-disciplinary. The attacker is expected to be familiar with commonly used bioinformatic tools and databases, such as BLAST and UniProt [20]. While deep cybersecurity knowledge is unnecessary, we assume the attacker is an open-minded enthusiast, influenced by cybersecurity literature on code obfuscation and classic encrypted computer viruses.

The required biological skills and resources are less demanding than those needed for recreating a viral pathogen, which, according to the World Health Organization (WHO), “did not require exceptional biochemical knowledge or skills, significant funds or significant time” [60, pp. 29-30]. The assumed attackers lack the expertise to invent new toxins not found in nature or databases—they are not innovators. They should have undergraduate-level hands-on experience in a biological laboratory, including skills in cell culture maintenance, transfection techniques, the use of restriction enzymes for DNA manipulation or splicing, and fundamental microbiology practices like bacterial transformation with cloned DNA and protein expression. In general, the attackers aim to minimize expenditures by avoiding costly equipment such as a desktop DNA synthesizer and sophisticated biological protocols (e.g., Gibson assembly to produce synthetic DNA from oligos) that demand significant hands-on experience.

The adversary should have an account with any synthetic DNA provider and $400-$2,000 to purchase synthetic DNA. They should have unrestricted access to laboratory equipment at home, work, or one of the many DIY lab spaces [7]. For example, one could purchase a Genetic Engineering Lab Kit for less than $1700.^6^ The total cost of the necessary equipment and consumables is approximately $5,000, including a Co2 Incubator and biological hood.

### 3.2 Choosing a toxin

This paper does not intend to provide guidelines for producing biological agents for bioweapons. However, it highlights key considerations for screening tools when assessing potential threats. Attackers are likely to select a toxin that is (1) sufficiently dangerous for their intended purpose, (2) practical to handle and deliver, and (3) feasible to produce in cells they can grow. Among toxins meeting these criteria, attackers would prioritize those that (4) can be obfuscated most effectively.

### 3.3 Below the DNA screening radar

Toxins are obfuscated by splitting into fragments, which are then interleaved with benign DNA.

The primary weakness addressed in this paper is overlooking potentially dangerous DNA sequences shorter than 50 bp (see § 2.2). Obfuscation enables a toxin to evade detection by protocols implemented under the HHS guidelines. Next, we elaborate on obfuscations based on splicing and restriction enzymes. Specific details of the biological protocols of both obfuscation methods will not be provided out of caution since the biosecurity community does not currently have a clear consensus on what laboratory details should be considered a potential information hazard [52]. Accordingly, for this presentation, we take a conservative position and note only that the methods are readily accessible to most laboratory biologists (see ethics considerations).

In addition to the two obfuscation methods elaborated in this paper, we have previously considered [8] an obfuscation based on CRISPR — a well-known modern DNA editing technique. Here, we do not discuss this obfuscation method, despite intriguing past results, because it requires a higher level of biological sophistication to implement [21, 50].

#### 3.3.1 Splicing-based obfuscation

DNA obfuscation based on splicing (Algorithm 1) is straight-forward and stealthy, but the reconstruction is not guaranteed, and its effectiveness depends on the specific introns and host cells.

##### Sequence design

Given a toxin selected by the attacker, Algorithm 1 outlines the main steps for designing an obfuscated DNA encoding this toxin. This obfuscation method relies on the splicing machinery most commonly found in eukaryotic cells, though it is also present in some bacteria. First, the attacker constructs a toxin-encoding DNA sequence using codons optimized for their preferred cell cultures (line 1). Although all codons can be used to express proteins in most cells, using optimized codons generally improves the efficiency of protein expression [73]. Next, the toxin-encoding DNA is split into *k* evenly sized fragments (line 2), where *k* is determined by the attacker (discussed in § 6). Finally, in lines 3-6, the toxin fragments are interleaved with an intron. The intron used for obfuscation is selected based on the splicing machinery available in the attacker’s preferred cell cultures. The resulting sequence is flanked by a promoter and a terminator to facilitate transcription.

###### Algorithm 1 Splicing-based toxin obfuscation

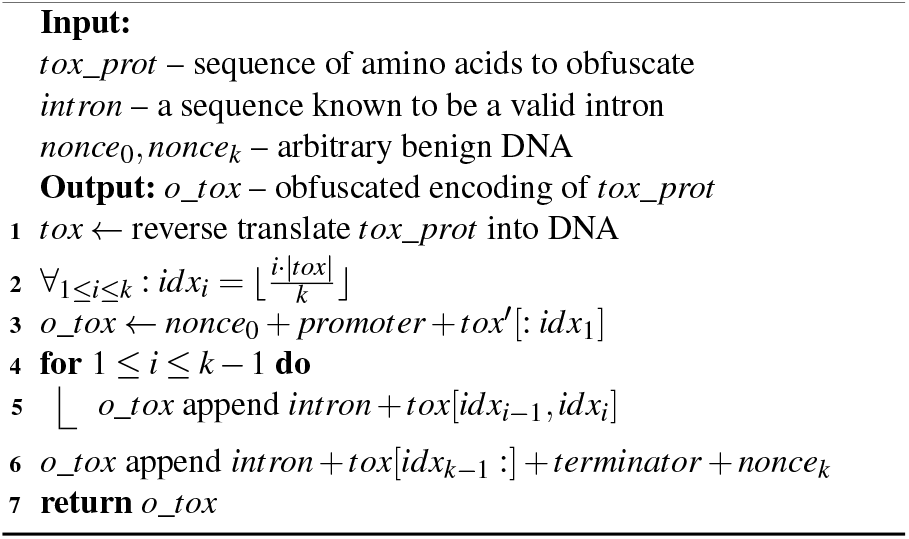

We assume that benign genes, denoted as *nonce*_0_ and *nonce*_*k*_, are appended to the beginning and end of the obfuscated DNA. These arbitrary benign DNA sequences are unlikely to be inserted between the toxin fragments (exons) because intron sequences affect their folding and, consequently, the splicing process. Therefore, an intermediate-level attacker would likely use well-known introns.

##### Reconstruction protocol

Reconstructing the spliced toxin is as straightforward as standard protein expression. The protocol would include: (1) transfection of the DNA obtained from the synthetic DNA provider using one of the standard techniques, and (2) growing the cells. Although splicing can also be performed in vitro the attacker is more likely to choose eukaryotic cells to leverage the splicing machinery and protein expression. This is because splicing is a multi-step biological process that is technically challenging to reproduce in vitro, while in vivo, it occurs naturally in the correct context.

#### 3.3.2 Restriction-based obfuscation

DNA obfuscation based on restriction enzymes involves a more complex design stage (Algorithm 2) but a more controllable reconstruction phase with a higher likelihood of success. ***Sequence design***. Restriction-based obfuscation is similar to splicing-based obfuscation but is subject to additional constraints imposed by restriction enzymes. Most notably, the toxin encoding DNA cannot be split into small fragments arbitrarily, but rather only at specific restriction sites. The location and the number of restriction sites depend on the DNA coding of the toxic protein. Non-optimal coding may have a better distribution of restriction sites, increasing the efficacy of the obfuscation while decreasing the efficacy of the protein expression compared to the optimal coding. This trade-off incites an interesting combinatorial optimization problem, which is out of the scope of this article. Here, we assume a brute force approach where the attacker considers all DNA codings of the toxic protein with at most *d* non-optimal codons (line 1).

The core of the obfuscation algorithm (lines 2-5) is finding the best set of *k* − 1 restriction enzymes that would cut the toxin encoding DNA into *k* most evenly sized fragments. The trade-off associated with the number of fragments is discussed in § 6. Algorithm 2 searches for each restriction site within each DNA variant (*tox* ∈ *D*, line 2) and records those restriction enzymes whose cleavage sites appear exactly once in *tox*. If a restriction enzyme cleaves *tox* in more than one location, then, during the reconstruction phase, it will create many disconnected pieces of DNA that may ligate and recombine in an uncontrolled manner. In the likely scenario of more than *k*− 1 different restriction sites in a DNA variant, the attacker will inspect all combinations (line 2). In lines 4-5, the attacker selects a combination of restriction enzymes that results in the most evenly sized fragments. Since all combinations result in exactly *k* fragments, the harmonic mean of their lengths is maximized when they are evenly sized.

Finally (line 6), toxin fragments are interleaved with an arbitrary benign nonce to separate the malicious fragments.

The promoter and terminator are inserted immediately before the first fragment and after the last fragment, respectively. Every cleavage point between consecutive toxin fragments 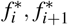 is transformed into two cleavage points: (1) between 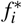 and *nonce*_*i*_, where the end of 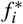 consists of the restriction site, and (2) between *nonce*_*i*_ and 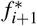,where the respective restriction site is prepended to 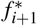.This will enable two cuts on both sides of the nonce using the same restriction enzyme, allowing the two fragments to ligate.

##### Example 1

*Consider, for example, the obfuscation process depicted in Figure 2. The toxin is 105bp long and the attacker splits it into two fragments (k* = 2*). The restriction site of AgeI is roughly in the middle of the sequence* 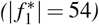.*The toxin is cut into two fragments (0-54) and (55-105), interleaved with benign genes represented by PuroR as well as Luciferase and IRES2. This layout of benign genes is a common practice in biology that may distract an inattentive analyst. The small blue bars near the scissors indicate the AgeI restriction sites. The first AgeI site is part of the end of the first fragment (N-Toxin), while the second is prepended to the second fragment (C-Toxin)*.

**Figure 2:**
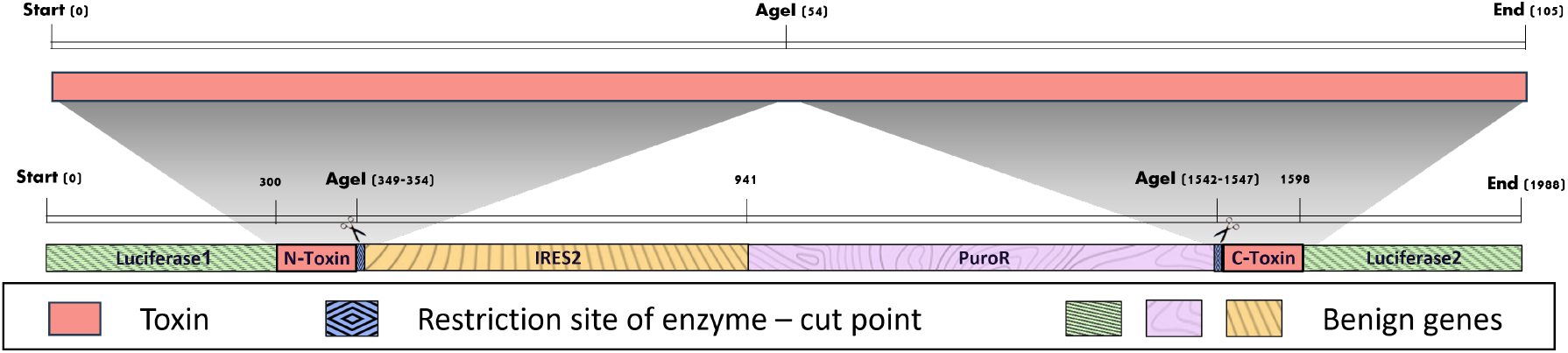
Example of a malicious DNA order based on a toxin, obfuscated by a restriction enzyme.

#### Reconstruction protocol

The reconstruction of the obfuscated toxin can be performed by iteratively applying the restriction enzymes. Both the splitting algorithm and reconstruction approach were validated *in vitro* using standard laboratory methods and a non-controlled sequence as the target for reconstruction. (see, for example, the many restriction digest protocols on https://openwetware.org/).

## 4 Gene Edit Distance (GED)

To harden synthetic DNA order screening against splitting-based obfuscation, we propose Gene Edit Distance (GED) designed to assess the difficulty of assembling a toxin from a DNA sequence instead of just checking if the DNA contains a toxin.

The GED algorithm scans the query sequence to find all fragments of a toxin. Then, the GED score quantifies the effort needed to assemble the toxin from these fragments using a series of cuts and repairs. Although designed with a biosecurity focus, GED can quantify the effort required to assemble any target sequence *t* from a query sequence *q*.

### Algorithm 2: Restriction enzyme obfuscation

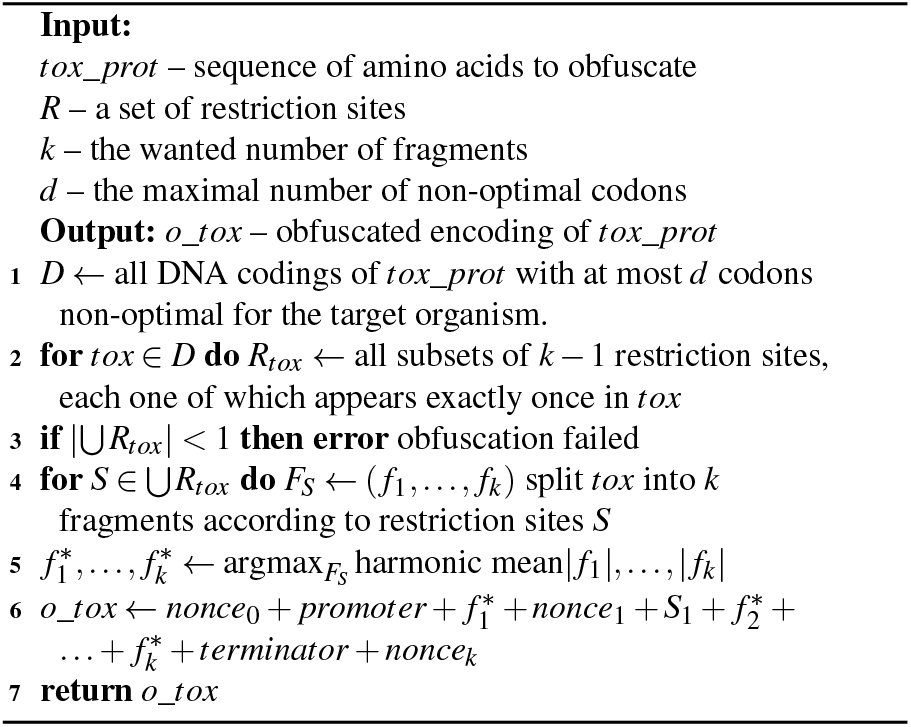

### 4.1 Sequence alignment preliminaries

Sequence alignment, a pivotal process in bioinformatics and biosecurity, involves arranging and comparing nucleotide or amino acid sequences to identify similarities and differences. This method is essential for understanding the functional and structural aspects of biological molecules, aiding in tasks such as uncovering evolutionary relationships and predicting gene or protein function. The Basic Local Alignment Search Tool (BLAST) [5] was one of the earliest and most influential frameworks developed for sequence alignment in bioinformatics. BLAST algorithms are designed to efficiently perform rapid searches in large sequence databases while maintaining a balance between speed and alignment accuracy. In BLAST, a **hit** represents a sequence in a database that contains at least one segment similar to the query sequence. The matching segments within the query and target sequences, exhibiting a high degree of similarity, are called SP. Each High Scoring Pair (HSP) is assigned a score, reflecting the degree of similarity between the respective sequences.

Next, we introduce the necessary principles and notations of sequence alignment. Let *q* denote a **query sequence** and *t* denote a **target sequence**. Let *q*[*i*] denote the *i*’th character in *q*. BLAST operates by matching n-grams (short sequences of letters) and extending these matches to form local **alignments** between the sequences.

Let 𝒜_*q,t*_ be a set of such alignments between *q* and *t* found by BLAST. Every alignment in α ∈ 𝒜_*q,t*_ is represented as a set of index pairs (*i, j*), where *i* corresponds to position in the query *q* and *j* to a position in the target sequence *t*. Gaps are represented by (*i*, ⊥) or (⊥, *j*), where ⊥ means that the argument character is not aligned. Alignments are monotonic, ensuring that if *i*_1_ < *i*_2_, then *j*_1_ < *j*_2_.

The score of an alignment α is computed based on the number of **matched** characters (*M* = | {(*i, j*) ∈ α : *q*[*i*] = *t*[*j*]}|); the number of **mismatched** characters (*MM* = |{(*i, j*) α ∈: *q*[*i*] ≠ *t*[*j*]); the number of **gap openings** in both the query and the target sequences and the total **gap extentions**, as follows. A **gap opening** refers to the first unaligned character in a sequence, while a **gap extension** accounts for consecutive unaligned characters following a gap opening. Formally the number of gap openings is *G* = | {(⊥, *j*) ∈α ∧ (⊥, *j* −1) ∈ α |+| (*i*, ⊥) ∈ α ∧ (*i*− 1, ⊥) ∈ α| and the total extent of gaps is *GX* = |{(⊥, *j*) ∈ α ∧ (⊥, *j*− 1) ∈ α | +| (*i*, ⊥) ∈ α ∧ (*i*− 1, ⊥) ∈α|. There is a reward for every matching character (*rm*) and penalties for every mismatching character (*pmm*), gap openings (*pgo*), and gap extensions (*pgx*). The rewards and penalties are configurable. The score of an alignment is:

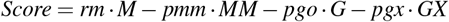

As demonstrated in Example 2, the alignment between sequences *q* and *t* results in matches, mismatches, gap openings, and a gap extension, which are reflected in the overall alignment score. To determine whether a *Score* of an alignment is significant, BLAST computes an **e-values**—the expected number of alignment with a similar *Score* that could occur by chance for a query of the same length in a database with the same number of entries. A lower e-value indicates a more statistically significant alignment.

#### Example 2

*Consider the following alignment:* α = {(1, 1), (2, 2), (3, ⊥), (4, 3), (⊥, 4), (⊥, 5), (5, 6), (6, 7)}.

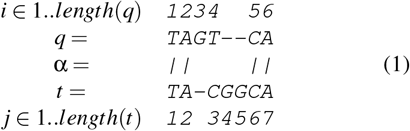

*The first two and last two characters are identical, the fourth is a mismatch, and there are two gap openings and one gap extension*.

### 4.2 The GED score

In standard biological sequence alignment, the typical objective is to identify regions of similarity between two or more sequences, which may indicate functional, structural, or evolutionary relationships. To achieve this, the match reward (*rm*), mismatch penalty (*pmm*), and gap penalties (*gpo, gpx*) must be well balanced when computing the alignment score. GED’s objective is more complex. On one hand, we need to identify short-matching regions with minimal gaps within the query sequence. On the other hand, we aim to concatenate these regions regardless of the gap length between them.

#### Example 3

*The default configuration for aligning two sequences with* BLAST *[6] is rm* = 2, *pmm* = 3, *pgo* = 5, *and pgx* = 2. *Consider an example, where* BLAST *returns three HSPs. As long as pgx* > 0, *these HSPs will not be merged by* BLAST *due to the length of the gap that would be opened in the target sequence. However, when removing a sequence between two consecutive toxin fragments using the restriction enzyme system, the distance between them is not critical*.

#### Gap removal penalty

To achieve GED’s objective, we introduce a new gap removal penalty (*prm*) that substitutes some gap opening and extension penalties in the target sequence.^7^

Let *g* = [*a, b*] be a gap in the target sequence 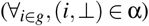 Removal of [*a, b*] from the query sequence requires two cuts, before *a* and after *b*, respectively, followed by cutting mechanisms (such as restriction enzymes or splicing). For more details on the process, refer to 3.3. These operations may fail. Let γ be the probability that a gap is successfully removed.

To define GED we ignore certain biological constraints, such as the required distance between two sites, assuming that cuts can occur at any point in a DNA sequence. This is a worst-case assumption overestimating the potential risk. Future versions of GED may take biological constraints into account to reduce false positives. Currently, in the absence of advanced adversarial techniques, GED detects obfuscated toxin with 100% accuracy, as shown in § 5.

If the gap *g* = [*a, b*] is removed, the alignment score will increase by *pgo* + *pgx* · (*b* − *a*) and decrease by *prm*. Let 𝒢_*q*_(α) and 𝒢_*t*_(α) be sets of all gaps in the query and target sequences, respectively, according to the alignment α. Let 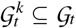 be a subset of the *k* longest gaps in the target sequence to be removed. The probability that all gaps in 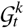 will be successfully removed is γ^*k*^. Next, we adjust the alignment score as a result of designating 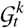 for removal.

##### Definition 1

**(Adjusted alignment score)** *Given an alignment* α; *a subset of gaps* 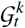 *in the target sequence of* α; *gap opening and gap extension penalties pgo and pgx respectively; gap removal penalty prm; and gap removal probability* γ; *we define the change in the alignment score as the result of the removal of the k longest gaps in the target sequence t as:*

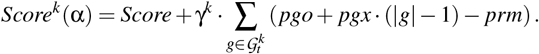

Next, we examine the choice of gaps to be removed (*G*_*trm*_) and the parametrization of *Score*^*k*^. We assume that the reward and penalties (*rm,pmm, pgo, pgx*) are set to their default values or according to biological considerations (which are beyond the scope of this article).

*Score*^*k*^ approaches *Score* when the number of gaps to be removed *k* increases and γ < 1. The choice of γ depends on the assumptions made regarding the expected biological effectiveness of the attacks given typical bioengineering tools used by potential victims today. γ = 0 signifies that no attacker could ever rely on the cutpoint mechanisms in the cells and the toxin obfuscation attack described in § 3.3 is impossible. γ = 1 signifies 100% success of gene editing, which leads to the successful decoding of toxin sequences regardless of the number of toxin fragments.

The gap removal penalty *prm* should be set to a value that justifies gap removal using bioengineering tools. For example, if the removal of gaps is only justified for gaps larger than a certain threshold size *x* bp, where gaps smaller than *x* are considered acceptable and do not require removal, then the gap removal penalty should be set to: *prm* = *pgo* + *pgx* · (*x* −1). Naturally, the longer a gap is, the more worthwhile its removal becomes. Thus, for the sake of computing *Score*^*k*^, only the longest gaps are selected. The number of gaps *k* to include in 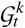 for the maximal *Score*^*k*^ depends on the parameters and can be selected using a grid search or simple hill-climbing algorithm.

##### Example 4

*Assume that we examine a 10000 bp query sequence, compared to a 2000 bp toxin. Assume a negatively scored optimal alignment between the two sequences that contains 2000 matching base pairs, a few mismatches, and 50 gaps with exponentially distributed lengths, as depicted in Figure 3a. Although such an alignment exists, it will never be returned by the* BLAST *search engine because its score is worse than random. Assuming default* BLAST *configuration, the specific alignment score could be lower than* − 4, 500. *Assume a gap removal penalty of prm* = 20. *Replacing gap opening and gap extension penalties with prm starting from the longest gaps would increase the score, as shown in Figure 3b. With* γ = 0.98, *setting k* = 10 *results in the highest Score*^*k*^ *value. With* γ = 0.99, *the adjusted alignment score can reach* 2, 000 *when k* = 13 *in this example*.

**Figure 3:**
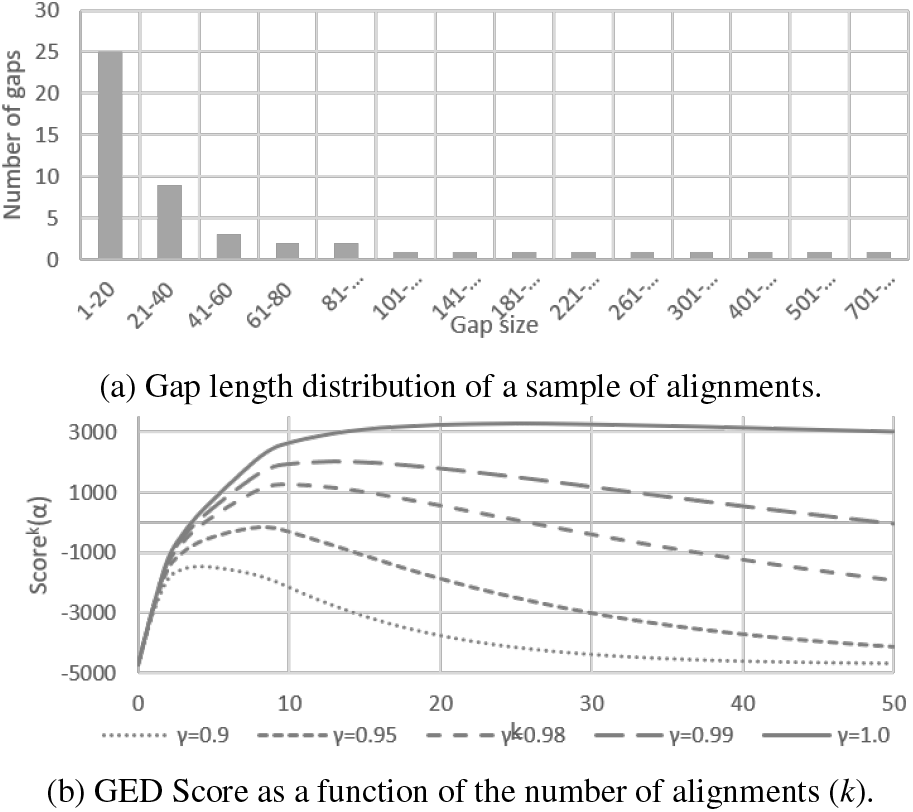
Example of alignment score correction using the GED Δ*Score*.

#### GED definition

Next, we define the gene edit distance as the optimal number of gaps (*k*) to remove in a query sequence such that the adjusted alignment score is maximized.

##### Definition 2

**(Unidirectional gene edit distance)** *The Gene Edit Distance (GED) from q to t is:*

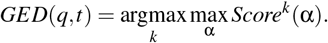

According to this definition *Score*^*GED*(*q,t*)^(α) has the maximal adjusted score. GED quantifies the effort required to transform *q* into *t* but not vice versa. As an academic exercise, one can define a true gene edit metric,^8^ but such a definition is beyond the scope of this security article.

### 4.3 The GED algorithm

Current BLAST implementations are optimized to penalize gaps, with a stronger focus on minimizing smaller gaps, which can make alignments with longer gaps less favorable under default settings. Thus, in order to reduce time-to-market and maintain backward compatibility with existing BLAST engines, we implement a GED heuristic as postprocessing of standard BLAST outputs. The pseudocode for GED is provided in Algorithm 3. Additionally, Appendix A illustrates the schematic flow of the algorithm and Appendix B demonstrates the application of GED to arbitrary strings.

#### Algorithm 3: Gene Edit Distance

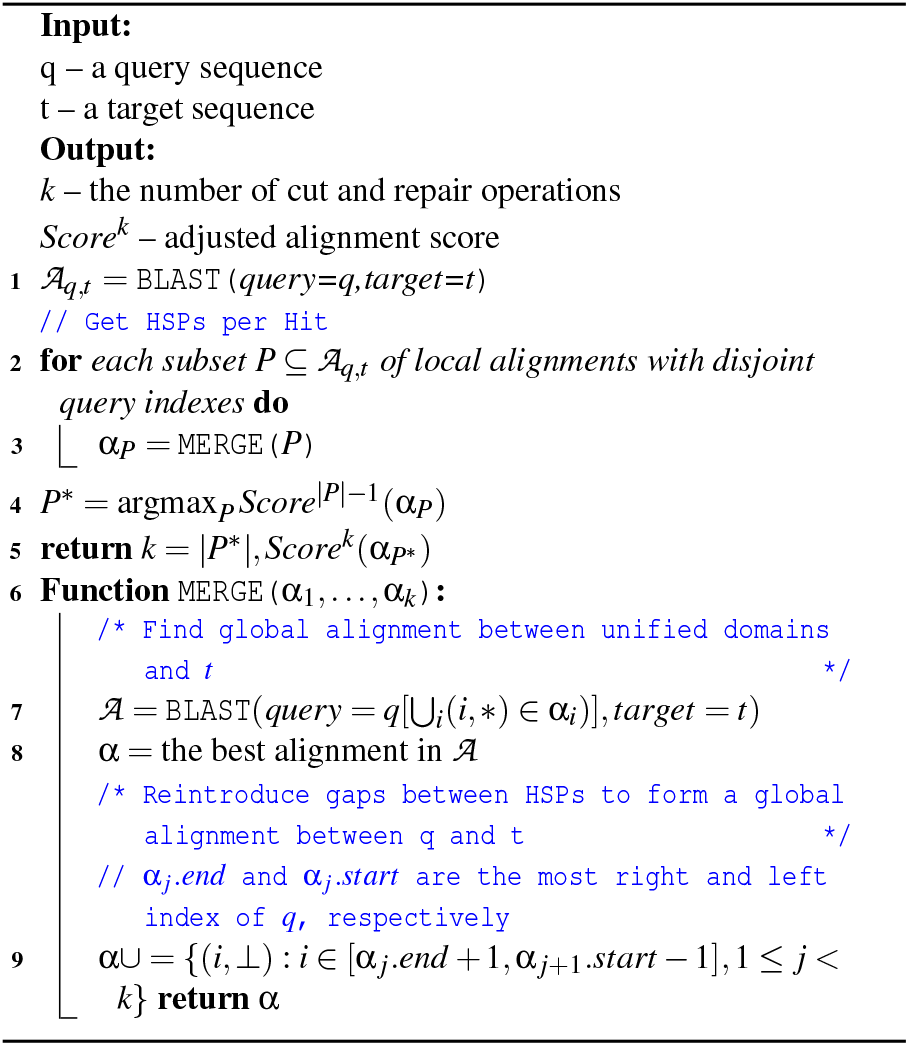

Standard BLAST algorithms compute a large set of small local alignments and extend these alignments when doing so increases the alignment score. We take a similar approach when computing a set of HSPs using standard BLAST configuration and merging them to maximize *Score*^*k*^. Recall that 𝒜_*q,t*_ is a set of local alignments between *q* and *t* returned by BLAST. Let .*start* and .*end* denote the first and the last position respectively, in the domain or the image of an alignment α ∈ 𝒜_*q,t*_ (Algorithm 3, line 1). We can use BLAST() to align the unified domains of α_1_, …, α_*k*_ ∈ 𝒜_*q,t*_ and the target sequence, i.e., when sequence fragments between the alignment domains are removed.

Let *P* = α_1_, …, α_*k*_ be a set of local alignments whose query indexes are disjoint, α_*i*_.*end* < α_*i*+1_.*start* (lines 2 and 3). The algorithm then selects the subset *P*^*^ that maximizes the adjusted alignment score *Score*^|*P*|−1^ (line 4). This subset *P*^*^ corresponds to the optimal set of disjoint alignments that, when merged, give the best alignment between the query and the target sequences (lines 7 and 8).

The output of Algorithm 3 is the number of cut and repair actions (i.e. gaps) required to reconstruct the target sequence from the query sequence and the adjusted alignment score. It’s important to highlight that GED’s versatility extends beyond the common scenario of two cutpoints. The algorithm is not limited and can effectively handle multiple cutpoints, broadening its applicability and utility in various biological contexts. This capability enhances its potential for accurately analyzing complex genetic data and reinforces its value in biological research and security measures.

To assess the risk posed by a query sequence, we compute its GED from all regulated sequences. The final judgment is based on the maximal adjusted score of the query sequence vs. any toxin sequence. This approach enables the identification of seemingly benign sequences that can be easily transformed into malicious sequences capable of producing dangerous products. Furthermore, this method is more resilient to attacks where a public gene database is poisoned with legitimate sequences. However, if legitimate sequences are maliciously marked as toxins, this could lead to false hits, requiring human review.

## 5 Experiments

In this section, we evaluate the impact of splitting-based DNA obfuscation on the alerts raised by biosecurity screening tools for a benchmark of toxins and venoms. Specifically, we address the following research questions:

- Does splitting-based obfuscation cause biosecurity screening to fail to detect controlled DNA sequences?
- Does detection of obfuscated sequences come at a cost of significant false positives?

### 5.1 Pre-screening and benchmark data

To answer these questions, we prepared a test dataset comprising obfuscated and non-obfuscated toxins, plus a collection of non-toxin sequences for evaluation of false positives.

#### Toxin dataset

Most DNA screening tools maintain their own databases, access to which is typically restricted due to a combination of biosecurity concerns and proprietary information. Therefore, we used an open toxic protein dataset sourced from UniProt’s Tox-Prot project [44], which we combined with prescreening of non-obfuscated sequences via the commercial biosecurity screening tools to identify 1045 sequences that should be flagged as sequences of concern potentially subject to regulatory control. BLAST, SeqScreen, and ToxinPred3 were not used for this purpose because these tools assess only structure and/or function, rather than the regulatory status of sequences. Nearly all of these sequences (1038 of 1045) are conotoxins, i.e., venoms from marine snails of the genus *Conus*, with the remaining 7 sequences being venoms from other marine snails and a species of sea anemone, flagged based on their similarity to conotoxins. Although a few sequences were missed by commercial Tool 2, as shown in the first row of Table 1, the tool provider confirmed that these sequences should have been flagged (false negatives not due to obfuscation).

**Table 1:** The fraction of sequences flagged as a potential toxin.

| True class | Obf. | n | BLAST short | BLAST non-short | GED | Tool 1 | Tool 2 | SeqScreen | ToxinPred3 |
| --- | --- | --- | --- | --- | --- | --- | --- | --- | --- |
| Toxin | — | 1045 | 100% | 99.7% | 100% | 100% | 99.6% | 97.6% | 95.1% |
|  | Intron | 1045 | 98.3 % | 95.8% | 100% | 90.6% | 88.8% | 87.6% | 0% |
|  | AfIII | 339 | 100% | 100% | 100% | 100% | 100% | 97.9% | 9.1% |
|  | Apal | 273 | 99.6 % | 99.6% | 100% | 98.5% | 98.2% | 94.5% | 33% |
|  | BamHI | 163 | 100% | 100% | 100% | 99.4% | 99.4% | 96.3% | 21.5% |
|  | EcoRI | 54 | 100% | 100% | 100% | 100% | 100% | 90.7% | 14.8% |
|  | EcoRV | 78 | 100% | 98.7% | 100% | 100% | 98.7% | 96.2% | 23.1% |
|  | HindIII | 581 | 100% | 100% | 100% | 99.8% | 99.8% | 96.9% | 14.1% |
|  | KpnI | 288 | 100% | 100% | 100% | 99.7% | 99% | 93.1% | 10.4% |
|  | NheI | 460 | 100% | 100% | 100% | 99.6% | 99.3% | 95.7% | 12% |
|  | NotI | 166 | 100% | 100% | 100% | 98.2% | 95.8% | 94.0% | 13.3% |
|  | XhoI | 306 | 100% | 100% | 100% | 97.7% | 96.7% | 93.5% | 15.4% |
| Non-toxin |  | 5571 | 0.5% | 0% | 0% | <1%* | <1%* | 0.04% | 6.6%** |

\* Non-toxin FP rates for Tool 1 and Tool 2 are the typical rates of total threat false positives (i.e., toxin, viral, viroidal, bacterial, and fungal) for these tools for real-world deployment on customer order streams, as reported by their maintainers.
\*\* Non-toxin FP rates for ToxinPred3 is derived from measurements reported in [64].

The amino acid sequences were then reverse-translated into DNA using standard codons, resulting in a collection of sequences ranging from 39 to 744 bp, with an average length of just 195 bp. Note that these sequences are, in general, significantly shorter than other controlled toxins, and much shorter than most genes from viral threats and virulence factors from bacterial and fungal threats. As such, this test set represents a particularly challenging scenario for testing obfuscation, and we may expect that tools will perform as well or better against obfuscation of other classes of threats besides the ones tested here.

#### Generation of obfuscated sequences

The 1045 controlled toxin sequences were obfuscated by inserting introns in the middle of each sequence according to Algorithm 1 with *k* = 2, using the commonly used beta-globin intron [69]. 7% of sequences obfuscated with introns had fragment sizes below 50 bp. The average length of an obfuscated sequence created with this method was 1,464 bp (roughly $300 at a cost of 0.20$/bp).

Consensus toxin sequences were additionally obfuscated with restriction enzymes according to Algorithm 2 with *k* = 2, using the 10 different restriction enzymes listed in the second column of Table 1 rows 3-13 (AfIll–XhoI). The prevalence of restriction sites varies among the toxin sequences: for example, HindIII was found in 581 sequences, while EcoRI was found only in 54, and 96 of the 1045 sequences did not contain any of the 10 restriction sites. In total there are 2,708 toxin sequences obfuscated with restriction enzymes. 16% of the 2,708 obfuscated sequences had roughly equal-sized fragments, each spanning 45%-55% of the toxin. 37 had the largest fragment shorter than 50 bp. The average length of an obfuscated sequence created with this method was 2,105 bp (roughly $420 at a cost of 0.20$/bp)

#### Non-toxin sequences

To evaluate false positive rates, we considered 5,571 presumably non-toxic protein sequences obtained from the ToxinPred database [64]. Pre-screening with commercial tools identified many sequences from controlled viral or cellular organisms in this dataset. Thus, we used it only to estimate false positives for GED, BLAST variants, and SeqScreen.

### 5.2 Hyper-parameters and setup

#### BLAST and GED

We evaluate whether obfuscation disrupts sequence alignment by running BLAST against a database consisting of only the 1045 controlled sequences. For this purpose, we used the blastn nucleotide sequence alignment instantiation of BLAST, executing with the default settings for short and non-short sequences. For BLAST short, we selected an e-value of 1E-5, allowing at most 0.5% FP. In contrast, the performance of the non-short variant was largely insensitive to variations in the e-value, therefore, the default setting was retained.

GED uses BLAST as indicated in the Algorithm 3, setting the e-value threshold to 0.01 to retrieve many HSPs. We set the GED score threshold to 35, which is the average value between the highest benign sequence to the lowest unobfuscated toxin sequence score. Obfuscated sequences were not used to set the threshold.

Finally, note that this experiment tests only whether a match can be made between an obfuscated sequence and its known non-obfuscated original, and the results produced are thus not a full biosecurity evaluation.

#### Commercial tools

We report the performance of two commercial DNA screening tools, whose names are replaced with pseudonyms Tool 1 and Tool 2 for purposes of confidentiality. The tools were operated by their respective maintainers.

#### SeqScreen and ToxinPred3

Both SeqScreen [4] and Toxin-Pred3 [64] utilize their own databases for analysis. SeqScreen was configured to run in sensitive windowed mode, using a window length of 200bp and an overlap of 100bp. To further enhance SeqScreen sensitivity, the query sequence was scanned with overlapping windows – a mode introduced in SeqScreen version 4.4 and updated in version 4.5, following the disclosure of preliminary results with obfuscated sequences. SeqScreen outputs multiple flags that indicate the function of the input protein. In this study, we consider only the *cytotoxicity* or *disable organ* flags. Note that SeqScreen flags the viral IRES gene, which we use as a benign nonce. We ignore this flag because there is no virus-related DNA among the 1045 consensus sequences. We deployed ToxinPred3 distributions with default settings on a Slurm cluster and configured it to detect at least 92% of unobfuscated toxins as reported in [64].

### 5.3 Screening performance

Screening results for all tools are presented in Table 1. Percentage values are obtained as the fraction of flagged sequences out of the total (*n*) in the respective row, except cells marked with an asterisk. The first 12 rows quantify the true positive (TP) rates—a fraction of flagged out all the toxin-encoding sequences. While the bottom row quantifies the false positive (FP) rate—a fraction of flagged out all the benign sequences. The FP rates of the commercial tools and ToxinPred are reported based on their own respective benchmarks.

#### Obfuscation does not disrupt alignment

Except in a small number of cases, BLAST alignment is not disrupted by obfuscation. This is particularly the case when running BLAST optimized for short alignment, as is expected given that some of the sequences are very short. Also as expected, BLAST does not find any significant alignments between toxin sequences and unrelated non-toxin sequences that are not in the database of sequences to align.

#### GED scoring separates obfuscated toxins and non-toxin genes

As expected, GED improves on BLAST alignment by considering how separated fragments could be merged via gene edits. Figure 4 provides further insight on this improvement by showing the score distributions for GED and both BLAST variants. Both BLAST short and non-short show a tradeoff between TP and FP manifested in the area under their receiver operating characteristic curve (AUC) of 0.98 and 0.95 respectively. GED provided a perfect separation between non-toxin and obfuscated toxin sequences with an AUC of 1. The separation gap between non-toxin with the highest score (32) and obfuscated toxin with the lowest score (40) was eight (8).

**Figure 4:**
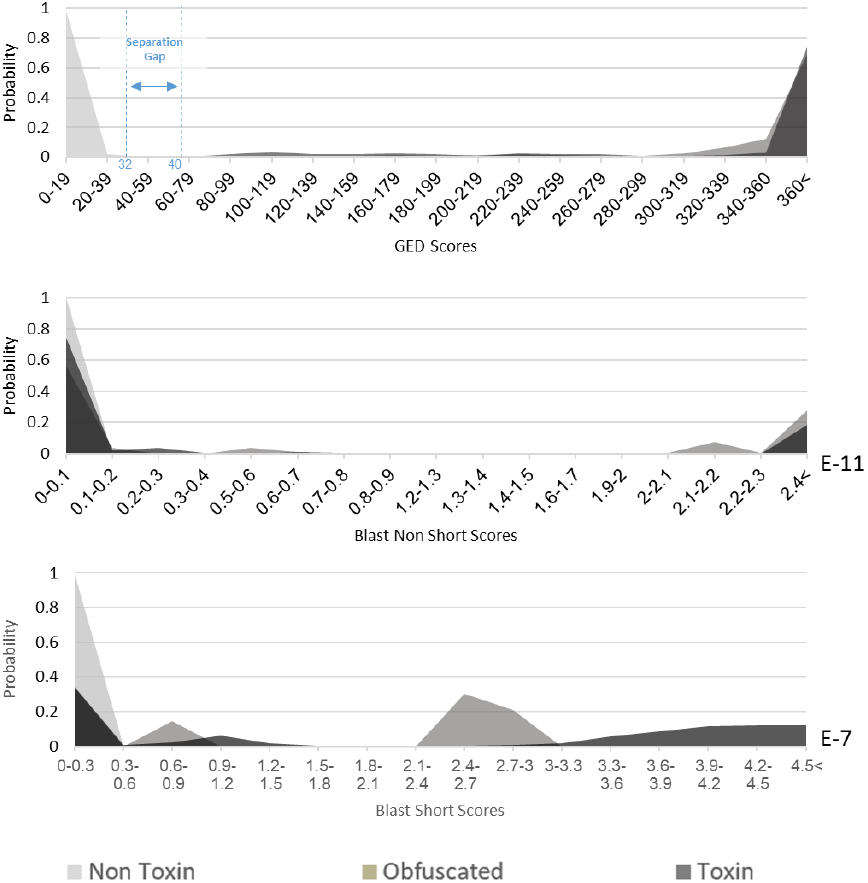
Distribution of GED scores compared with short and non-short BLAST, showing a separation gap in GED scores that is not present for either form of BLAST.

#### Commercial tools detect 50+ bp fragments reliably

The commercial screening tools perform a more complex analysis than BLAST or GED, as they have no *a priori* knowledge of the test sequences and are asking the general question of whether the sequences contain any potentially regulated sequences of concern. Overall, both commercial tools perform well against splitting-based obfuscation.

As evident from Table 1, however, some sequences may slip through biosecurity screening if they are obfuscated by being split into smaller fragments. This is expected, as both the 2023 US HHS Guidance and the 2024 Framework for Nucleic Acid Synthesis Screening target a minimum sequence length of 50 bp, and many of the sequences in the test set are less than 100 bp. Indeed, Table 2 shows that both tools are effective at detecting obfuscated toxins whenever at least one sequence fragment is at least 50 bp: Tool 1 misses only a single sequence (in three different obfuscated forms), and Tool 2 misses obfuscated sequences that it had also missed in non-obfuscated form. Both tools do flag many sequences shorter than 50 bp, but not all, and the likelihood of evading detection is thus higher for splicing-based obfuscation than for restriction-based obfuscation due to the balanced length of the fragments.

**Table 2:**
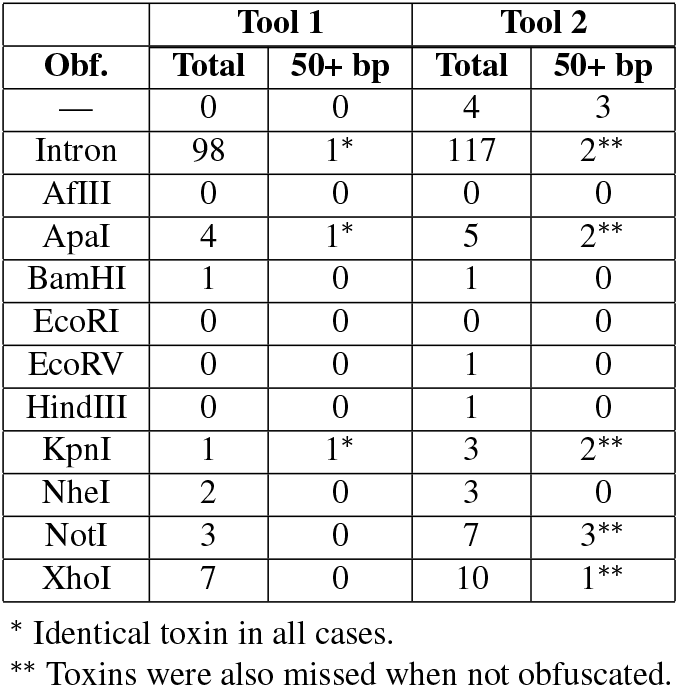
Sequences missed by commercial tools.

| Obf. | Tool 1 |  | Tool 2 |  |
| --- | --- | --- | --- | --- |
|  | Total | 50+ bp | Total | 50+ bp |
| — | 0 | 0 | 4 | 3 |
| Intron | 98 | 1* | 117 | 2** |
| AflIII | 0 | 0 | 0 | 0 |
| ApaI | 4 | 1* | 5 | 2** |
| BamHI | 1 | 0 | 1 | 0 |
| EcoRI | 0 | 0 | 0 | 0 |
| EcoRV | 0 | 0 | 1 | 0 |
| HindIII | 0 | 0 | 1 | 0 |
| KpnI | 1 | 1* | 3 | 2** |
| NheI | 2 | 0 | 3 | 0 |
| NotI | 3 | 0 | 7 | 3** |
| XhoI | 7 | 0 | 10 | 1** |

\* Identical toxin in all cases.
\*\* Toxins were also missed when not obfuscated.

#### ToxinPred and SeqScreen

Table 1 shows that SeqScreen catches nearly all non-obfuscated sequences, though slightly less than the commercial tools. SeqScreen also performs well against splitting-based obfuscation. As with the commercial tools, obfuscation does allow some sequences to slip through, with the impact of obfuscation being somewhat higher for SeqScreen than for the commercial tools. ToxinPred, on the other hand, is fairly effective at detecting non-obfuscated sequences, but performs poorly on obfuscated sequences.

### 5.4 Reconstruction proof of concept

This paragraph is limited in detail due to the ethical and biosecurity concerns as mentioned in § 3.3 and the ethics considerations section. Introns inserted into non-controlled DNA (§ 3.3.1) were correctly identified by multiple intron-detection software tools (SpliceRover,^9^ NNSPLICE,^10^, and NetGene2^11^). An experiment with an intron inserted within a green fluorescent protein [69] was successfully reproduced in a lab. DNA editing using restriction enzymes is ubiquitous in molecular biology [66], and there is no need to prove it works. Nevertheless, to validate the claimed threat model, students participating in this research have successfully reconstructed non-controlled DNA obfuscated according to § 3.3.2. The evidence reported here should not be considered a biological benchmark.

## 6 Discussion

We have presented and evaluated a novel category of potential vulnerabilities in DNA biosecurity screening arising from obfuscation techniques based on splitting sequences via splicing or restriction enzymes. Both methods have the potential to circumvent screening either by the addition of benign “distractor” material or by reducing the length of contiguous toxin-encoding DNA fragments below the 50 bp detection threshold recommended by the HHS Guidelines. This approach may seem like a straightforward obfuscation technique, but its biological implementation presents several challenges, as discussed below.

### 6.1 Conclusions from DNA obfuscation

Both restriction-based obfuscation and splicing-based obfuscation impose costs on an attacker. Restriction-based obfuscation requires a lower level of biological expertise since DNA restriction is ubiquitous in biological laboratories. However, obfuscation based on restriction enzymes requires selecting appropriate restriction enzymes and target sites, as shown in § 3.3.2 and Algorithm 2, and often leaves long fragments of toxin-encoding DNA detectable by screening tools. Designing splicing-based obfuscation is algorithmically simpler, as shown in § 3.3.1 and Algorithm 1, and the sequences are stealthier, as demonstrated by the results in § 5.3 and Table 1. Splicing mechanisms are much more complicated biologically, however, being restricted to certain classes of organisms and influenced by complex biological contexts that are not fully understood. Consequently, if splicing-based techniques fail, troubleshooting would be significantly more challenging for attackers compared to methods relying on restriction enzymes.

We attribute elevated false negatives of splicing-based obfuscation compared to restriction-based obfuscation to the production of evenly sized fragments with this technique, which reduces the length of the longest fragment. The attacker can also potentially reduce the size of toxin fragments to evade detection by increasing the number of fragments, either by inserting additional restriction or splice sites. Increasing the number of fragments thus increases the difficulty of detection, but also complicates the reconstruction of the attacker’s desired sequences. The *Best Match* principle suggests that between the fragments of malicious DNA, an attacker should insert benign genes with high homology to the malicious DNA. However, according to the feedback from screening tool providers, the type of genes present between the malicious fragments should not affect the screening results of Tool 1, Tool 2, or SeqScreen in windowed mode.

### 6.2 Implications for DNA screening

Our findings confirm that, as expected, commercial DNA screening tools detect obfuscated sequences very well, particularly those where at least one fragment is longer than 50 bp. Tool 1 missed a single sequence with a fragment longer than 50 bp, which can be attributed to the unlikely event of breaking a toxin characteristic signature at the restriction site. Experiencing this situation will lead to more robust signatures in the future. The toxins missed by Tool 2 further emphasize the need for a consistent common standard for defining sequences of concern and testing the ability of tools to detect such sequences [75].

Similarly, the SeqScreen patch addressed a weakness in its reporting procedures. While the required technology was already in place (allowing prompt patching), the missing outputs were discovered only after an adversarial examination of the tool. Finally, the case of ToxinPred3 demonstrates the difference between a scientific tool for analyzing sequence properties and systems designed for biosecurity screening.

To summarize, our red teaming resulted in two practical impacts: (1) discovering a weakness that led to the patching of a prominent open-source DNA screening tool (SeqScreen), and (2) advancing the ongoing development of community test sets. To further support the development of standards for detecting sequences of concern, we have contributed the benchmark dataset of obfuscated toxins to the test set development project currently led by the IGSC Test Set Working Group [75].

### 6.3 Adversarial analysis and future directions in biosecurity screening

This article has demonstrated how the application of cyber-security practices can help produce more robust, adversary-resilient biosecurity screening services, laying crucial groundwork for the eventual global regulation and enforcement of biosecurity screening. However, achieving this vision requires further advancements in DNA screening to address emerging threats and technological developments. First, susceptibility to sophisticated attacks remains a significant concern, particularly strategies involving the distribution and strategic combination of malicious DNA fragments across multiple orders. Although this approach poses a threat, it increases the biological sophistication level required to integrate disparate genetic material. The suggested GED metric and the principle of quantifying the effort to transform a seemingly benign sequence into a regulated (i.e., malicious) sequence is a potential step toward increasing the resilience of biosecurity screening, though additional testing and development is required to determine whether it is practical for inclusion in biosecurity systems.

One concern for future biosecurity screening is the potential use of non-natural synthetic bases or amino-acids [33, 48]. Non-natural bases extend beyond the standard A, T, C, and G nucleotides, creating what is called XNA molecules. Once synthetic DNA providers start selling custom XNA molecules, it will be imperative to update screening algorithms to account for the inclusion of these non-natural bases. Since there are no appropriate databases to retrieve the XNA function, future biosecurity screening must rely on artificial intelligence (AI) and structural features of XNA.

Advanced AI is also a major biosecurity concern. New unnatural proteins designed de-novo by AI algorithms to perform some biological function will likely be missed by screening tools based on signatures or similarity to known biological sequences [10, 37] (including GED). Future biosecurity developments may evaluate the protein function based on its 3D structure [77]. Finally, advanced AI agents based on large language models may potentially reduce the biological sophistication level required to produce dangerous agents despite biosecruity screening [23, 26]. Safeguards of large language models [25] must, therefore, consider the sensitivity of biological protocols that enable the construction of dangerous biological agents.

### 6.4 Summary

Most importantly, there is a general opportunity for improving biosecurity through increased cross-disciplinary collaboration of cybersecurity researchers with the biosecurity community.

For example, efficient, secure, and privacy-preserving threat information sharing in biosecurity is an important challenge [67] where cybersecurity threat intelligence practices [2] may be extremely valuable. While it is not yet clear which cybersecurity techniques will best generalize for the case of biosecurity, this study shows the positive benefits that can be accrued from even relatively simple collaborations, and it is reasonable to expect that many more such opportunities abound.

#### Biosafety

The research reported in this article, including biological proof of concept, was conducted in accordance with local biosafety regulations and with appropriate permits.

#### Ethics considerations

This paper highlights a critical aspect of biosecurity, particularly in a landscape where the synthesis of hazardous DNA poses a potential threat. The threat model outlined in § 3.1 specifically addresses the risk of low-profile bioterrorists attempting to exploit vulnerabilities in the synthetic DNA screening process. The defense-evasion technique was disclosed to the IGSC consortium during their monthly meeting. Interested parties participated in a thorough vulnerability evaluation. Part of the results of this evaluation are reported in this article. Vulnerabilities found in commercial tools were acknowledged and deemed as not violating biosecurity regulations except in one case that may require deeper investigation. A weakness arising from improper use of the open-source SeqScreen software was acknowledged and promptly patched by the developers in SeqScreen v4.4 with database v23.3. We also informed the developers of the ToxinPred tool about the identified issue.

Before submission, this article was sent for community review for potential information hazards to a set of stakeholders in government, industry, and academia, and certain details of methods were abstracted or omitted based on community feedback; we thank these reviewers for their input. Abstracting and omitting some details on the biological protocols also raises the entry barrier for novice do-it-yourself biologists to play with dangerous substances. Details of the biological protocols kept in the manuscript are sufficient for an experienced biologist to assess their correctness and reproduce the experiments. USENIX Security reviewers can find the omitted details in the accompanying letter, alongside revision criteria and a list of changes made to the paper.

To address potential biosecurity concerns, we released a narrowly focused dataset consisting of the long toxins that are detected by existing screening tools. Similarly, not to make the DNA obfuscation accessible for script-kiddies, we refrain from publishing the implementation of the obfuscation algorithms. Instead, we provided a pseudo-code representation sufficient to convey the core concepts and technical ideas of the obfuscation process. These measures were taken to balance transparency with responsibility, ensuring that our findings advance the field while minimizing potential risks.

## Acknowledgments

Blinded. Thanks to Bob Weinberg and Xiaoping Sun for contributing specific sequences and materials for laboratory validation.^12^ This document does not contain technology or technical data controlled under either U.S. International Traffic in Arms Regulation or U.S. Export Administration Regulations. The views and conclusions contained herein are those of the authors and should not be interpreted as necessarily representing the official policies or endorsements, either expressed or implied, of the ODNI, IARPA, ARO, or the U.S. Government.

## Open Science

GED algorithm is available in the following public repository: https://anonymous.4open.science/r/gene-edit-distance-C5B6/

Which contains:

- Source code for the GED algorithm.
- Detailed documentation for setting up and running the algorithm.
- Operating instructions.
- A sample dataset in FASTA format, containing 54 sequences derived from 10 toxins. The dataset includes 10 sequences obfuscated by introns and others modified by various restriction enzymes, providing a robust testbed to evaluate the GED algorithm.

## A GED workflow

The diagram in Figure 5 illustrates Algorithm 3, with each stage in the diagram corresponding to the numbered lines in the algorithm.

**Figure 5:**
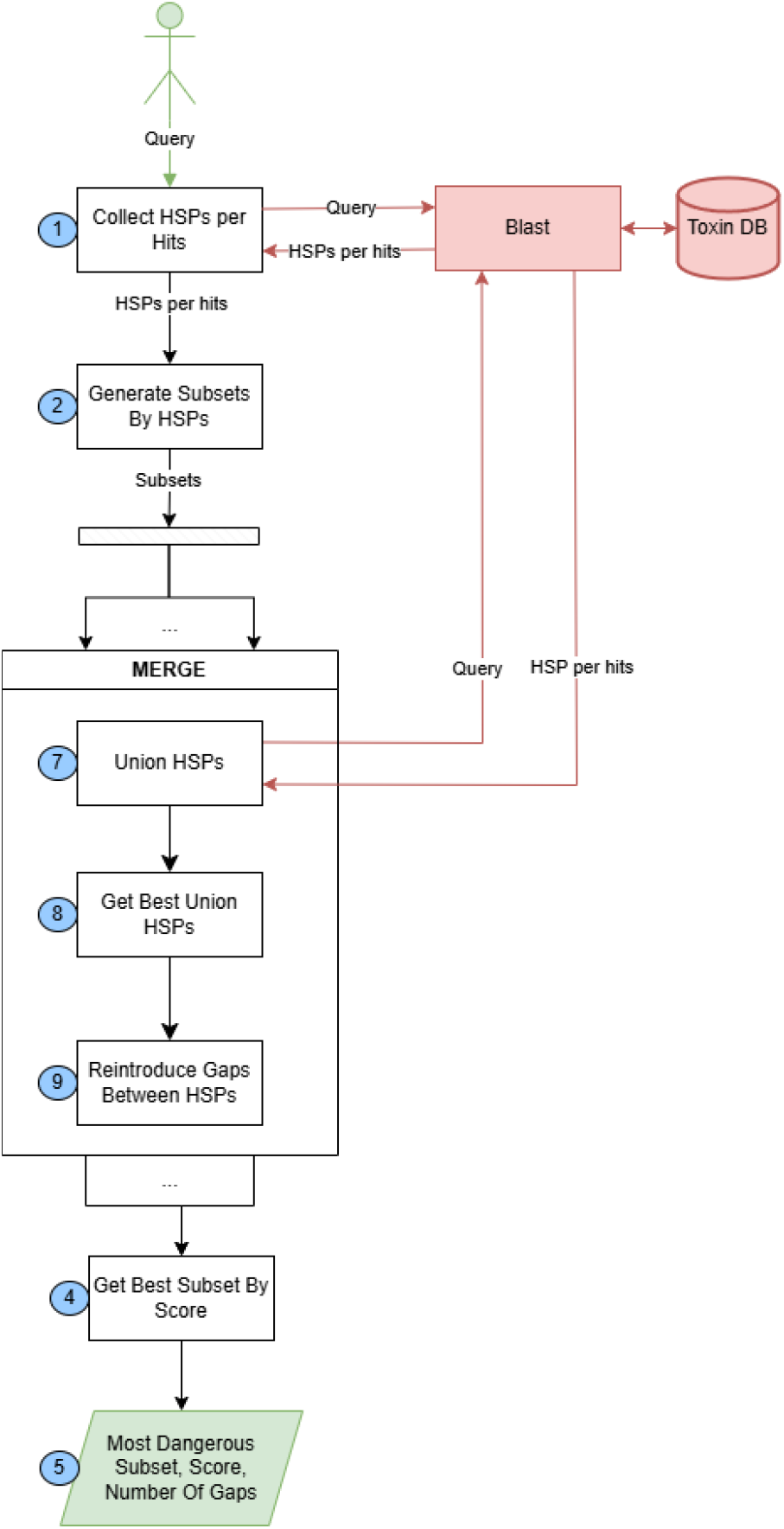
GED Workflow

## B GED example

To better illustrate the algorithm, we provide an example. Assume that the algorithm takes the following arbitrary string as input: AppleBananaOrangeKiwiCherryMango. Assume that in step (1) Collect Hits, we identified the following partially matching sequence: BananaKiwiMango, with the corresponding HSPs: [(5, 11), (17, 21), (27, 32)].

The second step (2) Generate Subset HSPs results in: [[(5, 11)], [(17, 21)], [(27, 32)], [(5, 11), (17, 21)], [(5, 11), (27, 32)], [(17, 21), (27, 32)], [(5, 11), (17, 21), (27, 32)]).

For each subset, the HSPs are merged in step (7) Union HSPs to form the unified domains: [[BananaKiwi], [BananaMango], [KiwiMango], [BananaKiwiMango]].

Subsequently, these unions are submitted to BLAST to obtain local alignment scores. For this example, let’s assume the results are as follows (query, new HSP, score):

- (BananaKiwi, score = 46)
- (BananaMango, score = 29)
- (KiwiMango, score = 43)
- (BananaKiwiMango, score = 55) Next, in step (9) gaps are reintroduced between the HSPs:
- Banana− − − − −−Kiwi
- Banana− − − − − − − − − − − − − − −−Mango
- Kiwi− − − − −−Mango
- Banana− − − − −−Kiwi− − − − −−Mango

Then, the scores are calculated, and the algorithm decides whether to calculate gaps or cut the gaps based on the BLAST parameters (match reward and penalties for mismatches, gap opening, gap extension, and gap removal). Assuming the parameters are set to: rm = 1, pmm = -2, pgo = 5, pgx = 2, and prm = 16. The scores for each case would be as follows:

- Banana − − − − −− Kiwi, score = max(46 - 1 * pgo - 5 * pgx, 46 - prm) = > 31
- Banana − − − − − − − − − − − − − − −− Mango, score = max(29 - 1 * pgo - 15 * pgx, 29 - prm) = > 13
- Kiwi − − − − −− Mango, score = max(43 - 1 * pgo - 5 * pgx, 43 - prm) = > 28
- Banana − − − − −− − − − − −− Kiwi Mango, score = max(75 - 2 * pgo - 12 * pgx, 55 - 1 * pgo - 6 * pgx - prm) = > 21

Finally, in step (4), the algorithm selects the best subset with the highest score. In this example, the optimal subset is Banana− − − − −−Kiwi.

## Footnotes

1 Types of RNA molecules and the specifics of transcription machinery are not relevant for this discussion.

2 Regulation effective from October 2026, but already implemented by the commercial tools in this study.

3 https://securedna.org/

4 https://ibbis.bio/our-work/common-mechanism/

5 https://www.aclid.bio/

6 https://www.the-odin.com/genetic-engineering-home-lab-kit/

7 Note that variation of gap penalties for query and target sequences has successfully been used in the past for other use cases [71].

8 Metrics are symmetric.

9 http://bioit2.irc.ugent.be/rover/splicerover

10 https://www.fruitfly.org/seq_tools/splice.html

11 https://services.healthtech.dtu.dk/services/NetGene2-2.42/

12 To reviewers: This acknowledgment does not violate the blind submission policy. It is a CC attribution required for the genetic materials we used.

